# The Molecular Subtyping Resource (MouSR): a user-friendly tool for rapid biological discovery from human or mouse transcriptional data

**DOI:** 10.1101/2021.08.12.456127

**Authors:** Baharak Ahmaderaghi, Raheleh Amirkhah, James Jackson, Tamsin RM Lannagan, Kathryn Gilroy, Sudhir B Malla, Keara L Redmond, ACRCelerate Consortium, Tim Maughan, Simon Leedham, Andrew S Campbell, Owen J Sansom, Mark Lawler, Philip D Dunne

## Abstract

Generation of transcriptional data has dramatically increased in the last decade, driving the development of analytical algorithms that enable interrogation of the biology underpinning the profiled samples. However, these resources require users to have expertise in data wrangling and analytics, reducing opportunities for biological discovery by “wet-lab” users with a limited programming skillset. Although commercial solutions exist, costs for software access can be prohibitive for academic research groups.

To address these challenges, we have developed an open source and user-friendly data analysis platform for on-the-fly bioinformatic interrogation of transcriptional data derived from human or mouse tissue, called “MouSR”. This internet-accessible analytical tool, https://mousr.qub.ac.uk/, enables users to easily interrogate their data using an intuitive “point and click” interface, which includes a suite of molecular characterisation options including QC, differential gene expression, gene set enrichment and microenvironmental cell population analyses from RNA-Seq. Users are provided with adjustable options for analysis parameters to generate results that can be saved as publication-quality images. To highlight its ability to perform high quality data analysis, we utilise the MouSR tool to interrogate our recently published tumour dataset, derived from genetically engineered mouse models and matched organoids, where we rapidly reproduced the key transcriptional findings.

The MouSR online tool provides a unique freely-available option for users to perform rapid transcriptomic analyses and comprehensive interrogation of the signalling underpinning transcriptional datasets, which alleviates a major bottleneck for biological discovery.

## INTRODUCTION

In the years since the first whole genome was sequenced, the costs associated with the generation of molecular “big data” have decreased rapidly, to a point where the data handling, rather than data generation, is the limiting factor in large biological discovery programmes. Furthermore, large repositories (such as the TCGA (NCI and National Human Genome Research Institute, 2006) and Gene Expression Omnibus (Edgar et al., 2002)), now provide free access to publicly-available molecular data. Large international molecular subtyping projects have markedly improved our biological understanding of cancer (Sohn et al., 2017), but in doing so they have created a critical bottleneck in terms of data reduction, analysis and interpretation, resulting in an urgent need for solutions that enable rapid biological interrogation of large datasets (Cerami et al., 2012).

Given the relative paucity of translational bioinformaticians within many research groups (Gao et al., 2013), there is a need for wet-lab researchers to have access to user-friendly analytic platforms that provide rapid and statistically controlled algorithms to perform common transcriptional analysis tasks, alongside an array of tools for visualising and interrogating the resulting data. For these tools to be widely adopted, they will need to provide the non-computational user with intuitive “point and click” options for transcriptional analyses, rather than programming-based options. To address this need, we have developed the molecular subtyping resource; MouSR; https://mousr.qub.ac.uk/, which enables individual non-computational end-users to pursue lines of investigation on transcriptional data within their domain of interest/area of expertise without the need for expertise in scripting. The MouSR platform enables interrogation of existing publicly-available or in-house transcriptional data and analytics from either human or mouse models, within a standardised molecular stratification environment.

## Methods

### Access and Requirements

The MouSR App was built using R (v3.2) and is running on the shiny server (v1.5.16) hosted on the Queen’s University Belfast virtual server CentOS 7, 64-bit, Intel Xeon Gold 6130 CPU @2.10 GHZ, 16 Core. The service was given extra security and protection by being placed behind a proxy service, which meant the server itself is never directly exposed to the internet. This configuration may be of benefit, where possible, to other potential users who wish to install their own MouSR version. The system is accessible via all web browsers tested, and on both Linux and Windows systems, via https://mousr.qub.ac.uk/. However, Chrome, Edge and Firefox are the recommended browsers for the best App experience.

### File formats

#### Importing Text Files (CSV or TXT)

This App accepts two commonly used text file formats. For “.csv” comma separated values (Comma delimited), data files are converted to text files, with each line corresponding to a row and all the fields in each line separated by commas, whereas in “.txt” (Tab delimited) all the fields in each line are typically separated by tabs. The uploaded files size is limited to 30 MB.

#### Transposing data tool for import

The Transcriptional Data Matrix file to be uploaded in the App has a defined format to follow, however for those users where their file is not in the same orientation as the suggested format in the App, a link to the online transposing tool has been embedded in the introduction section of the App and can be accessed via this link (Data Design Group, 2013).

### Tools embedded within the MouSR application

#### Principal Component Analysis (PCA)

The PCA analysis is performed by the prcomp function in the R stats package (Jolliffe and Cadima, 2016). PCA is defined by a transformation of a high dimensional vector space into a low dimensional space. It uses linear combinations of the original data to define a new set of variables that are referred to as principal components.

#### Multi-Dimensional Scaling Analysis (MDS)

We used the cmdscale function in the R stats package to perform MDS analysis (Young, 2013). Unlike the PCA method that minimises dimensions while preserving covariance of the data, MDS minimises dimensions and preserves distance between data points. However, both methods can provide similar results, if the covariance in data and Euclidean distance measure between data points in high dimension is equal. MDS uses the similarity matrix as input, which has an advantage over PCA as it can be applied directly to pairwise-compared banding patterns. The ‘% Variance’ describes how much of the total variance is explained by each of the components with respect to the whole (the sum). ‘% Variance’ values are shown on the axis labels.

#### Differential gene expression analysis (DESeq2)

The differential gene expression analysis is performed based on the negative binomial distribution using the DESeq2 R package (version 1.24.0) (Love et al, 2014). DESeq2 is a count-based statistical method that performs an internal normalization where estimated variance-mean is calculated for each gene across all samples. DESeq2 also estimates the gene-wise dispersion and logarithmic fold changes, a dispersion value is estimated for each gene through a model fit procedure, and differential expression is tested, based on a model using the negative binomial generalized linear distribution (Love et al, 2014). We used the DESeq2 package to normalize the data and identify genes, which are differentially expressed between the two main groups selected by the user.

#### Single sample Gene Set Enrichment Analysis classification (ssGSEA)

is performed using the GSVA package version 1.32.0. The R package msigdbr version 7.1.1 is also used to retrieve mouse/human Hallmark and biological processes (GO_BP) gene sets and applied to the samples (Hänzelmann et al., 2013).

#### Gene Set Enrichment Analysis (GSEA)

This method consists of three steps: Enrichment Score (ES) is calculated, reflecting the degree to which a set of genes is overrepresented at the top or bottom of the entire ranked list. Second, the statistical significance of the ES is estimated by using an empirical phenotype-based permutation test procedure that preserves the complex correlation structure of the gene expression data. Finally, after an entire database of gene sets is evaluated, the estimated significance level is adjusted to account for multiple hypothesis testing by first calculating Normalised Enrichment Score (NES), based on dividing the actual enrichment score by the mean of enrichment scores against all permutations of the dataset, then calculating the False Discovery Rate (FDR) corresponding to each NES. In this paper, GSEA is performed on log expression ratio using fgsea, an R package which is a fast implementation of pre-ranked GSEA (Sergushichev, 2019).

#### Microenvironment Cell Population counter (MCP)

The MCPcounter and Murine MCP (mMCP) counter R packages are used to estimate the quantity of several immune and stromal cell populations from heterogeneous transcriptomic data for human and murine samples, respectively (Petitprez et al., 2020) (Becht et al., 2016).

### Packages used

**S**hiny (Chang et al., 2018), shinythemes (Chang et al., 2021), shinydashboard (Chang et al., 2018), shinycustomloader (Tanaka and Niichan, 2018), shinycssloaders (Sali et al., 2020), ggplot2 (Wickham et al., 2020), tibble (Müller et al., 2021), DESeq2 (Love et al., 2014), limma (Smyth et.al, 2015), plyr (Wickham, 2020), biomaRt (Durinck et al., 2005), heatmaply (Galili et al., 2021), reshape (Wickham, 2018), plotly (Sievert, 2019), WGCNA (Horvath et al., 2021), lattice (Sarkar et al., 2020), pheatmap (Kolde, 2019), RColorBrewer (Neuwirth, 2014), GSVA (Hänzelmann et al., 2013), rlist (Ren, 2016), msigdbr (Dolgalev, 2020), tidyverse (Wickham, 2021), mMCPcounter (Petitprez et al., 2020), MCPcounter (Becht et al., 2016), magrittr (Bache et al., 2020), dplyr (Wickham et al., 2021), ggrepel (Slowikowski et al., 2021), readxl (Wickham et al., 2019), DT (Xie et al., 2021), colourpicker (Attali and Griswold, 2020), fgsea (Sergushichev, 2019), enrichplot (Yu and Hu, 2021).

### File outputs and modifiable formats

The customised options for plots that are common in all App sections are: turning labels on/off, justifying height and width of the plot and having different downloading format (png/svg). Additionally, some analytical panels have extra customised options in their filtering criteria, in order to provide an easier-to-use environment for the users, such as changing the colours, size of labels and points, assigning output filename, adding legend and changing scale.

### Data availability

The complete source code for the MouSR App can be launched on any system that has R/RStudio installed and is available to download at https://github.com/Dunne-Group/MouSR (Site will go live on publication acceptance). In addition, we are open to mutually-beneficial collaborations to further develop the MouSR analytic options.

If you use MouSR in your work, please remember to cite this paper.

## RESULTS

### MouSR Interface

The MouSR (https://mousr.qub.ac.uk/) platform is implemented as an open-source application that enables non-computational users to rapidly go from an existing data matrix (multiple formats) to biologically meaningful results in a user-friendly way. At each step of the process, users have the option to modify outputs, through a series of on-the-fly customisable graphics that can all be downloaded at high resolution for future use. The standard pipeline includes initial data QC assessments, followed by differential analyses, both single sample and group-wise gene set enrichment analyses and microenvironment population counters, producing publication-ready data (Figure 1). The system uses a species-agnostic “blind embedding” format, where users can upload data derived from any patient sample or *in vitro/in vivo* model as either ungrouped individual samples or multiple samples within experimental groups. The species-specific selection options available for downstream analyses in MouSR enable users to perform biological discovery/validation on transcriptional data derived from human or mouse origin.

**Figure 1:**
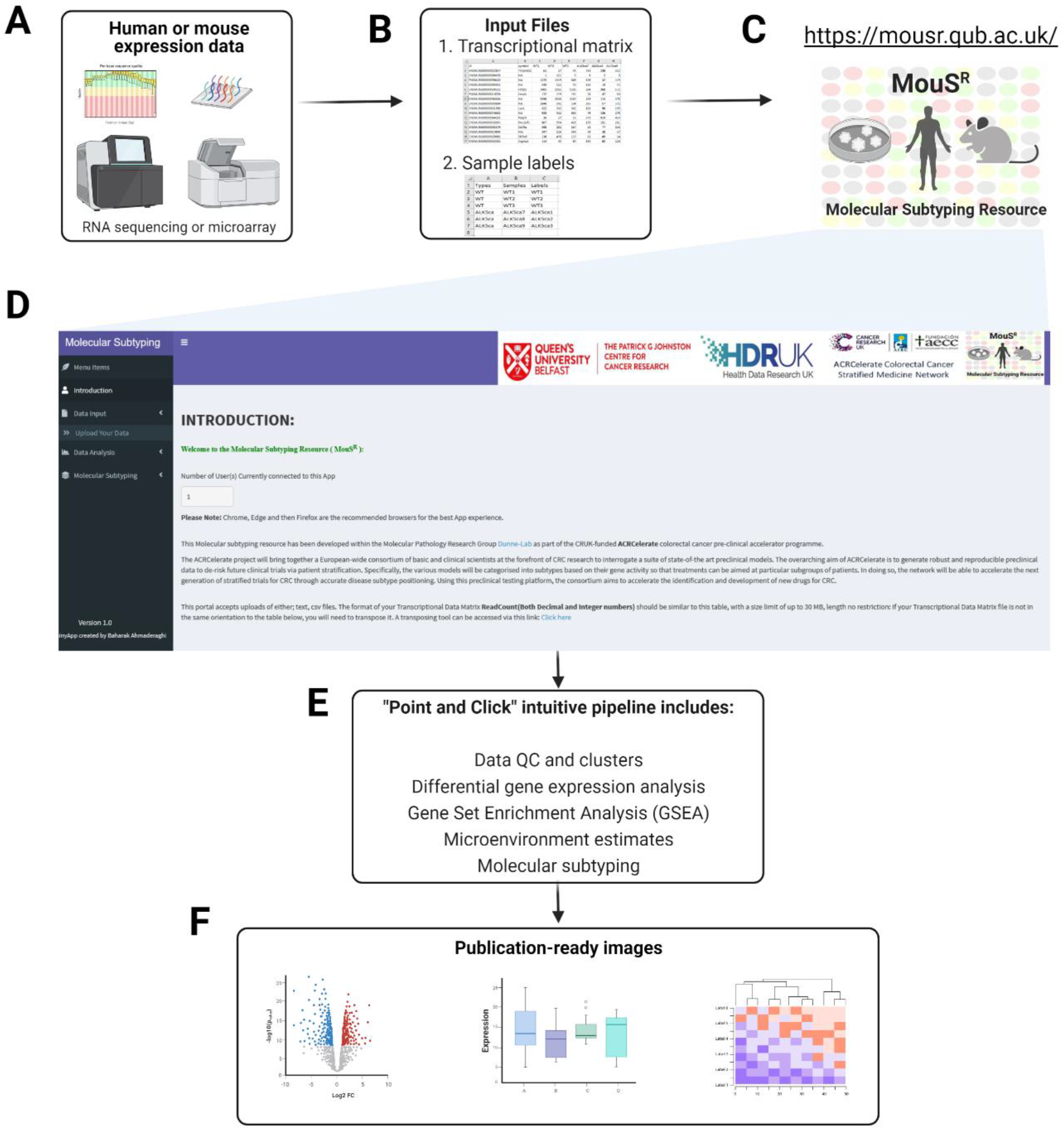
The MouSR workflow and outputs: Utilising transcriptional data derived from either human or mouse tissue/cells (A) users are required to have a transcriptional data matrix and sample information as the input for the MouSR pipeline, accessible via https://mousr.qub.ac.uk/ (B and C). From the introduction page (D), users upload their files and are then presented with a series of point-and-click options for initial data QC, differential analysis, molecular signaling and microenvironment characterisation (E) that can be saved as high-resolution image files for further use (F). As an example of the adaptable nature of the system at each stage, users have options for bespoke formatting, design and labelling of the resulting plots, which can all be downloaded and saved in a publication-ready format. Figure created using BioRender.

To highlight the functionality of the MouSR system, in this paper we will focus on colorectal cancer (CRC), using our previously published and comprehensively characterised genetically engineered mouse model tumour and organoid datasets, comprising of n=28 samples from matched tumour tissue (n=4 groups) and genotype-matched organoids (n=3 groups) derived from n=4 different genotypes (Jackstadt et al., 2019) (Jackstadt, 2019).

Accessing the MouSR website presents the user with a general introduction landing page, providing an overview of the application with instructions and exemplar formats that are required for use. The user interface structure has two main sections, namely 1) Data Input and 2) Data analysis, which will be described briefly, followed by an analysis of the CRC mouse exemplar dataset from Jackstadt and colleagues (Jackstadt et al., 2019) (Jackstadt, 2019) (Ahmaderaghi, 2021) and for convenience these data are included here as supplementary files (Supplementary files 1 and 2). In addition, a tutorial video is also includes to summarise the main features of the MouSR tool (Supplementary file 3).

### Data Input

The Data Input section is designed to have flexibility in terms of acceptable file/data formats, to enable users to upload their own data derived using a variety of transcriptional profiling platforms and normalisation procedures. Users are required to have two separate files; a transcriptional data matrix (input 1) and a sample information file (input 2) that will enable data analysis and generation of results (Figure 1).

In terms of data types for input 1, MouSR has successfully been tested using human and mouse data derived from a variety of microarray and RNAseq platforms and is adaptable enough to accept data that has been processed using a range of pipelines resulting in either integers (whole numbers, i.e. RNAseq read counts) or decimals (i.e. estimated read counts). However, as DESeq2 requires non-normalised counts as input, for users who select decimal input the differential expression options will not be accessible. Additionally, the MouSR system has been designed to accept the most common file formats; including csv, txt, and also to accept data files with various separators; including comma, semicolon or tabs. The data required for input 2 is a summary of basic information that relates to sample labels and groups.

Prior to uploading their data, users must ensure both files are in the recommended format described on the introduction page, which is aligned with a standard data matrix output containing gene ID and gene symbol columns followed by sample values and is in the format selected by the user according to their data types (default set as tab delimited/.txt file). To ensure that users with data in an orientation not supported by MouSR can use the tool, we have created a transpose link on the introduction page, which will adjust the transcriptional matrix using the Transpose CSV Tool (Data Design Group, 2013).

Once the files are in the correct format, the user is required to upload their two files into input 1 and input 2 (Figure 2A) using either a drag/drop from a folder, or by navigating to the file location using the browse function. When files are selected, a progress bar will immediately begin to indicate that the file is uploading until the upload is complete. By clicking on the submit button, the user interface becomes active, triggering the computational analysis on the background server, with input detail and input summary being displayed when complete, including information on number of samples, sample names, and number of expression values identified.

**Figure 2:**
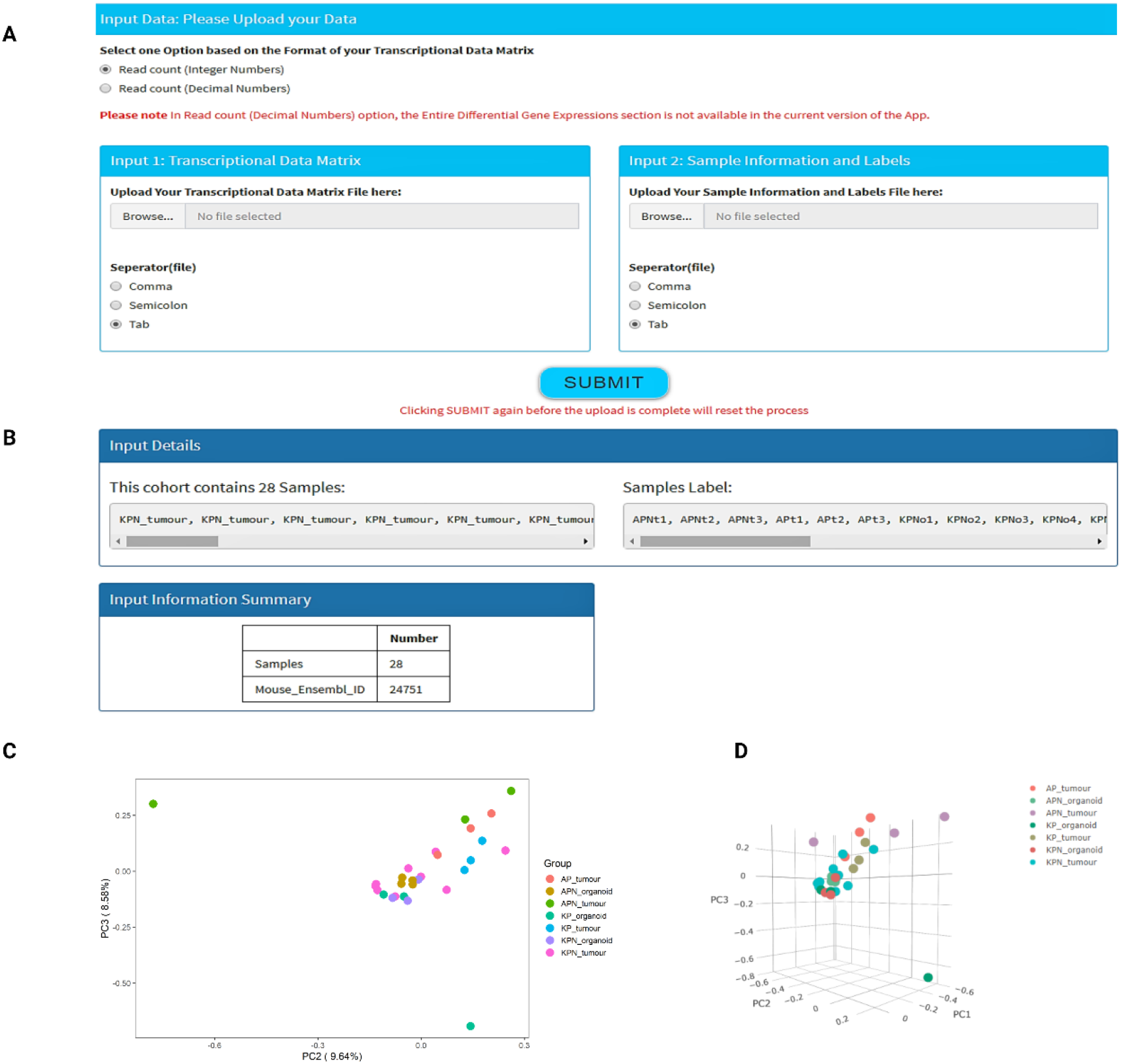
Data import and exploratory analysis. A) Two input files are required to begin the analytical pipeline in the App – a gene expression matrix and a meta data that includes sample group labelling. B) Following data upload, the data summary on samples will be displayed for quick review. C) Exploratory visualisation of data will be provided in a form of 2D PCA plot and D) 3D PCA plot with labelled sample groups.

Given the flexibility and the point-and-click design for downstream analyses, users must have their files in the correct format to proceed. At this stage, users can verify if their data are correctly loaded, or if modifications are required. Our exemplar files took <20 seconds from submission to display of input details, confirming that it consists of 24751 individual genes across 28 samples across 7 experimental groups (Figure 2B).

### Data Analysis

The *Data Analysis* section consists of four main subsections: 1) Principal Component Analysis (PCA) & Multi-Dimensional Scaling (MDS), 2) Differential Gene Expression Analysis (DGEA), 3) mouse and/or human Gene Set Enrichment Analysis (GSEA), and 4) mouse and/or human options Microenvironment Cell Population counter (mMCP/MCP-counter).

#### PCA and MDS

Principal Component Analysis (PCA) and Multi-Dimensional Scaling (MDS) are dimensionality-reduction methods that represent an initial step in assessing characteristics of any dataset (Jolliffe and Cadima, 2016) (Young, 2013). These options give the user an immediate overview of clustering of samples according to their experimental labels (described further in the Methods section).

##### PCA 2D/3D Plots

PCA plots enable the user to look at the principal components that describe the largest variability between samples in the dataset, where each data point corresponds to an individual sample. In MouSR users can also select any pair of the first three principal components for their static PCA plot and avail of customisable options, including the ability to choose a different colour for defined groups, turning labels on/off, justifying height and width of the plot, changing size of the labels/points, and having different downloading format (png/svg). In the 3D PCA plot option, MouSR exploits the functionality offered by the Plotly package (Sievert, 2019) to generate an interactive plot with adjustable features, giving users the option to rotate and zoom the graphic, isolate certain samples, alongside the ability to obtain sample information by hovering the mouse pointer over each data point. Instructions are displayed on the left-hand side of the plot, under the Plotly mode bar control, and hovering over the 3D PCA graphic itself will also reveal the built-in adjustable options above the sample labels on the top right. Furthermore, the colours of the data points are linked to the earlier 2D PCA colour option. Using our CRC mouse exemplar files (Jackstadt et al., 2019) (Jackstadt, 2019) (Ahmaderaghi, 2021), samples related to each experimental group are identifiable using the same colours in both the 2D (Figure 2C) and 3D options (Figure 2D).

##### MDS 2D Plot

The MDS plot has been added as a further option to project high dimensional data down to two dimensions, while preserving relative distances between observations (described in the Methods section). Again, the colour of the plot is linked to the 2D PCA.

### Differential Gene Expression Analysis

A primary objective of many gene expression experiments is to detect and analyse transcripts that display differential expression levels across different samples or experimental conditions. In MouSR, such analyses are made easy via a series of intuitive customisable options that enable selection of bespoke groups, thresholds, and filtering criteria.

#### Heatmap

MouSR, has been developed to ensure that a choice of different filtering options are provided. In the group comparison analysis panel, the comparison between two main groups is embedded from the labelling information the user uploaded in input 2, with the names of groups appearing as a list in dropdown menus for both group A and group B. This design provides the user with options to compare two individual groups, or to perform the comparison on up to 10 groups at each time, as users can pool up to 5 experimental groups in Group A versus up to 5 in Group B. The default Heatmap plot is generated based on log2FoldChange [-2,2] and an adjusted p-value cutoff of 0.05, using the “heatmaply” package (version 1.1.1) (Galili et al., 2021), however all of these options can be adjusted by the user (Figure 3A). Clicking the submit button will initiate the MouSR app to run DESeq2 on the data, producing customisable heatmaps, tables, boxplots, and volcano plots. Once complete, the heatmap plot details how many genes are either up- or down-regulated under these conditions. However, as with most features in MouSR, users have the option to adjust these to their own desired values, followed by clicking on “Update Plot” button to trigger the heatmap to be updated in real time. Users have the option to perform the clustering according to sample names or gene names or both, in order to visualize the differentially expressed genes. This interactive tool allows the inspection of a specific value by hovering the mouse over a cell, as well as zooming into a specific section of the figure by clicking and dragging around the relevant area. The differential gene expression data can also be displayed or downloaded as a table for analysis in other downstream tools, with samples as columns and gene names as rows. To demonstrate the utility of these features, we utilise the exemplar files to reproduce some of the main findings from the original Jackstadt et al. study (Jackstadt et al., 2019). Using MouSR, we first perform differential gene expression analysis comparing primary tumour data from four mouse genotypes (AP, APN vs KP, KPN), and plot the resulting differential genes using the heatmap tool (Fig 3B).

**Figure 3:**
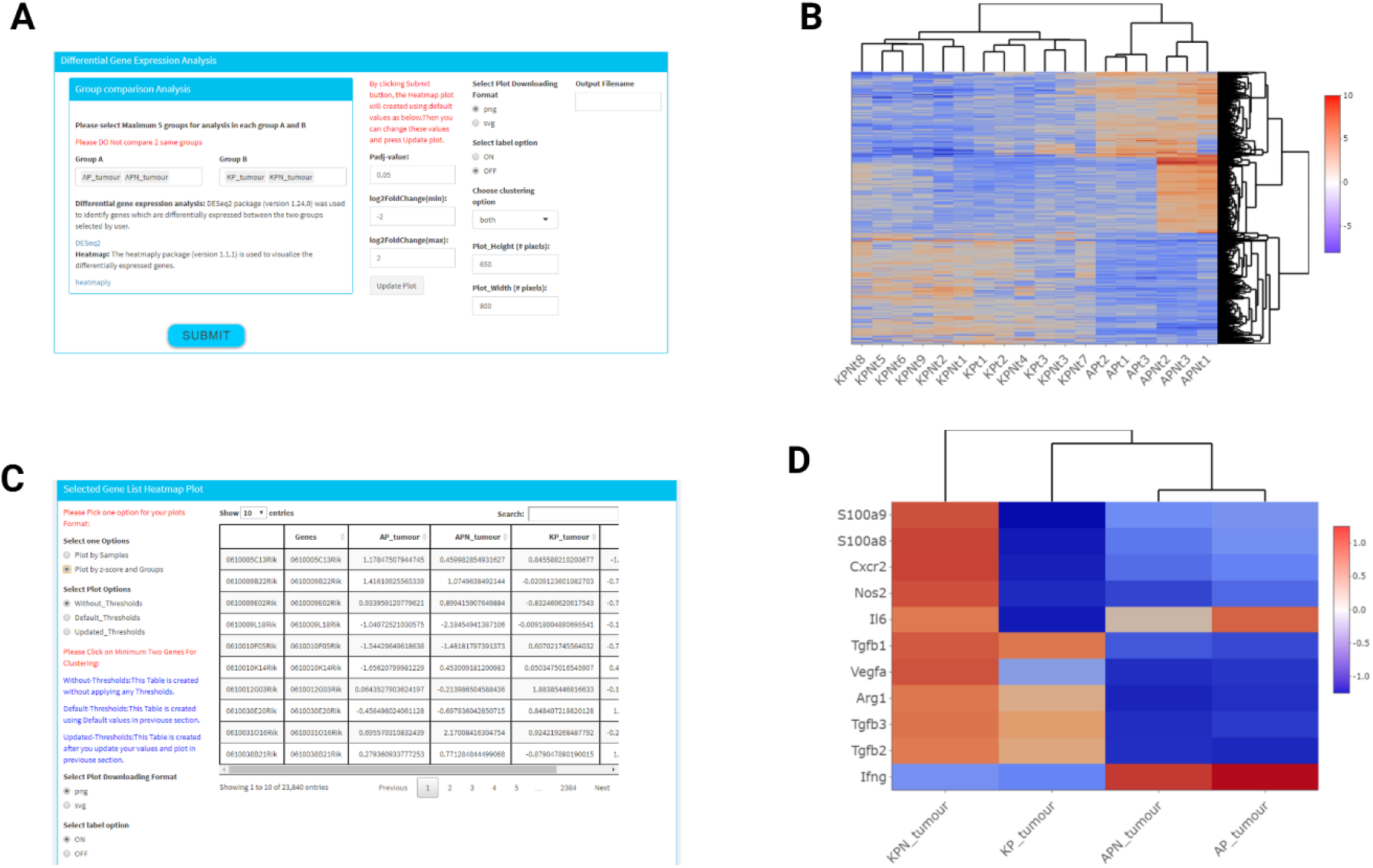
Differential gene expression analysis and visualisation options. A) Illustration of the various filtering options in the Heatmap panel for the gene expression in all the samples from the chosen groups. B) Heatmap depicts comparison between AP, APN vs KP, KPN across tumors. C) The selected gene side bar for various filtering options is visible to the left of the table. D) Heatmap indicates a reproducible result, compared to data from (Jackstadt et al., 2019) (Figure 6I) across the selected GEMM tumors.

#### Heatmap for selected Genes

This section provides the ability for users to create heatmaps for specific genes of interest (minimum of two genes) by clicking on the gene name in the table or by using the search bar on the top right corner of the table. Once selected, the heatmap will be created in real time as more genes are selected/deselected. Furthermore, there are two ways to create a heatmap, based on either individual sample values obtained during the differential analysis, or by creating an experimental group summary Z-score (scale between 1.25 and −1.25) according to each group analysed (Figure 3C). The table is created using three different options, the first one is for every gene uploaded in the original matrix, without applying any thresholds (Without_Thresholds), the second option (Default_Thresholds) is based on default values log2FoldChange [−2,2] and Padj-value: 0.05 in the previous section. The third option (Updated_Thresholds) will reflect any modifications to the default thresholds selected by the user during the previous differential step. From the original study, a number of specific markers were found to be differentially expressed between these models (Figure 6I in Jackstadt et al. (Jackstadt et al., 2019)), namely S100a9, S100a8, Cxcr2, Nos2, Il6, Tgfb1, Vegfa, Arg1, Tgfb3, Tgfb2 and Ifng. Assessment of gene expression levels for these individual genes produced a result in less than 30 seconds that was consistent with the original study, confirming the utility of the MouSR application (Figure 3D).

#### Volcano plot

Using the open-source tool VolcaNoseR (Luijsterburg and Goedhart, 2020) as inspiration, we have incorporated an interactive and customized volcano plot into MouSR using two different options. The first option is “using-plotly”, which exploits functions within the Plotly package (Sievert, 2019) that give the user the option to obtain essential information by hovering the mouse pointer over a dot showing the name of a corresponding gene. The second option, called “selectedGenes”, gives the users the option to annotate the volcano plot with up to 10 genes names (case-sensitive), which generates a new plot in real-time. As a default, the volcano plot shows the log2 of the fold change [−5,5] on the x-axis and minus log10 of the p-value on the y-axis, with a Significance threshold of 0.01 (Figure 4A). However, users have the option to adjust the size of data points, alongside options to modify parameters to their own desired values, followed by clicking on “Update Plot” button to trigger the volcano plot to be updated in real time. Using our exemplar dataset, we examined the transcriptome of KPN versus KP organoids, where we produce a volcano plot that demonstrates increased expression in Fjx1, Dtx1 and Tgfb2 (Figure 4B), similar to the published data (Figure 6A in Jackstadt et al. (Jackstadt et al., 2019)).

**Figure 4:**
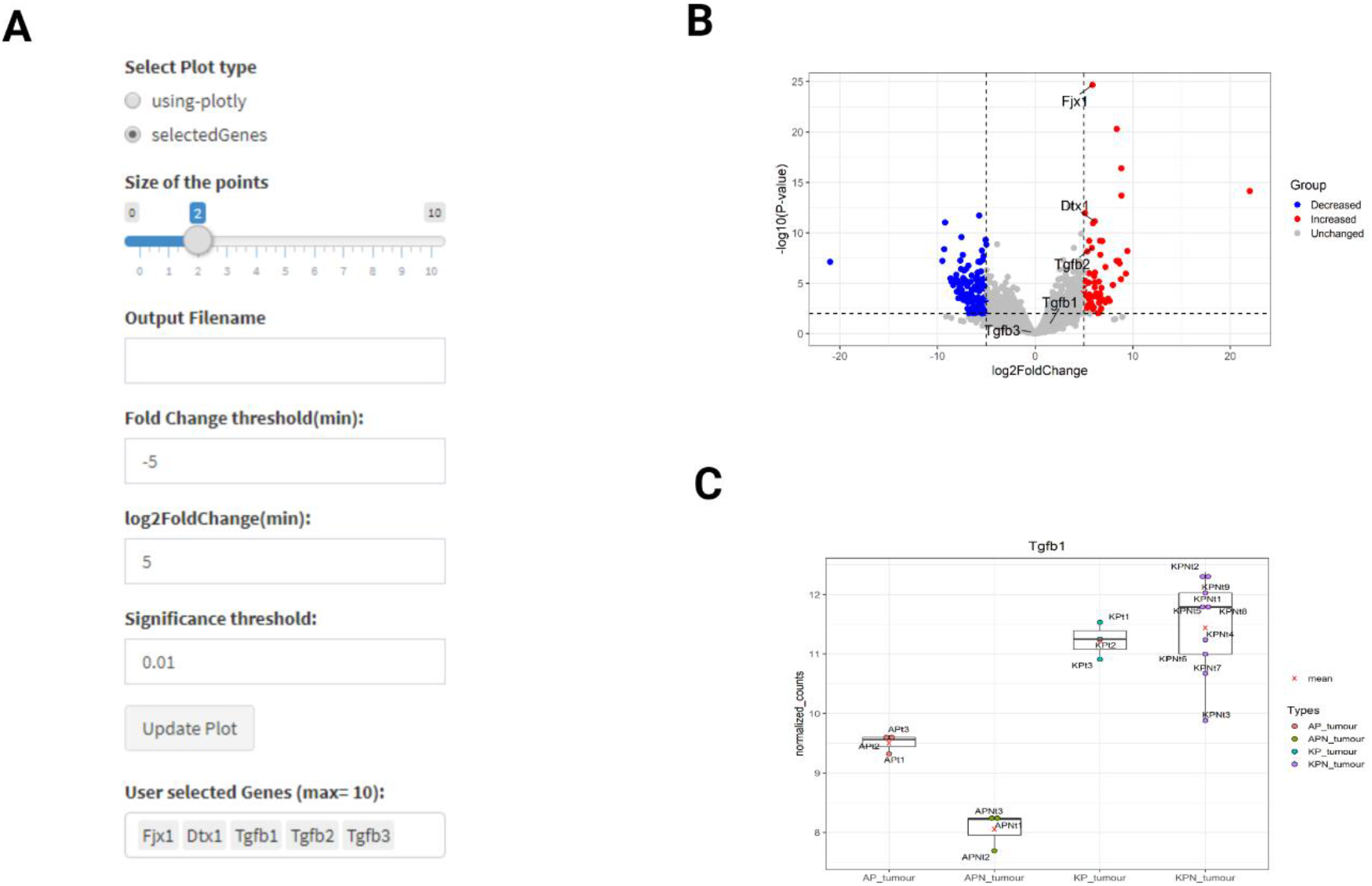
Gene Expression Levels and Volcano plot options. A) Volcano plot filtering options, by hovering over the plot using Plotly, the information related to each gene can be accessed immediately. B) Volcano plot displaying differentially expressed genes with highlighted key genes in text between KPN and KP organoids (reproducible result compared to data from (Jackstadt et al., 2019) Figure.6A). C) Boxplot displays normalised counts for Tgfb1 Expression compared between groups.

#### Gene Expression Levels

In this subsection, users can again select a specific gene name, via the table or the search option, to produce a boxplot of normalized count for expression values in each experimental group. In the original study, there was a focus on elevated Tgfb gene expression in the KP and KPN models compared to the AP and APN models. Again, using the intuitive MouSR system, we utilise the normalised counts for Tgfb1 expression plotted to reproduce this main finding (Figure 4C).

### Mouse/Human-specific Gene Set Enrichment Analysis classification (GSEA)

Since its introduction, GSEA (Mootha et al., 2003) (Subramanian, 2005) has become an essential part of the genomic analysis compendium of tools, due to its ability to measure and compare similarities or differences in experimentally validated biological signatures in transcriptional datasets. In MouSR, we provide options for both the original pairwise GSEA method (group A v B) and the modified single sample classification (ssGSEA), using the fgsea and GSVA packages (Hänzelmann et al., 2013) respectively, for both the Hallmark and Gene Ontology collections. Furthermore, in order to facilitate simultaneous classification between human- and mouse-derived data, we have extended our framework to provide an option for the users to choose between human or mouse analytical packages, based on their transcriptional data. Users also have the option of uploading their own bespoke list of genes or pathways of interest as an .rdata file. For users with a gene list from a spreadsheet, we have also created a side link that will convert a .txt file to .rdata, making this more user-friendly for non-computational users.

#### GSEA plot

In the GSEA plot section, users have the option to compare their two groups (selected during the differential analysis) with any specific gene sets within the Hallmark or Gene Ontology collections, which produces an enrichment plot and an indication of the number of leading-edge genes. For the Gene Ontology option, as the collection comprises over 7000 gene sets, only the first 50 pathways based on Enrichment Score (ES) will be available. The GSEA algorithm ranks genes based their expression, focussing on enrichment differences between samples belonging to two classes, labelled A or B.

### Mouse/Human-specific Microenvironment Cell Population counter (mMCP/MCP-counter)

The MCP algorithm gives an estimate of predefined immune and stromal cell populations from heterogeneous transcriptomic data (Becht et al., 2016). MouSR includes dual species templates to ensure users can assess either mouse or human data. For human, these populations include 8 immune populations (CD3+ T cells, CD8+ T cells, cytotoxic lymphocytes, NK cells, B lymphocytes, cells originating from monocytes (monocytic lineage), myeloid dendritic cells, neutrophils), and 2 stromal populations (endothelial cells and fibroblasts) (Becht et al., 2016). For mouse, these populations include 12 immune cell types (T cells, CD8^+^ T cells, NK cells, B-derived cells, memory B cells, monocytes/macrophages, monocytes, granulocytes, mast cells, eosinophils, neutrophils, and basophils) and 4 stromal populations (vessels, lymphatics, endothelial cells, and fibroblasts) (Petitprez et al., 2020).

Using the exemplar data, we perform GSEA using the Hallmarks collection on KP and KPN tumour samples, where, in line with the original publication, we observe an enrichment for TGF_BETA_SIGNALLING in KPN compared to KP tumour (Figure 5A). In addition to the pair-wise method, MouSR also enables users to perform single sample assessment using ssGSEA (Figure 5B) and MCP (Figure 5C) to assess enrichment in individual samples regardless of the experimental group. Given the adaptability of the MouSR tool, we will continue to add new options for data analyses, therefore features in a testing phase will be indicated as such (i.e. beta version).

**Figure 5:**
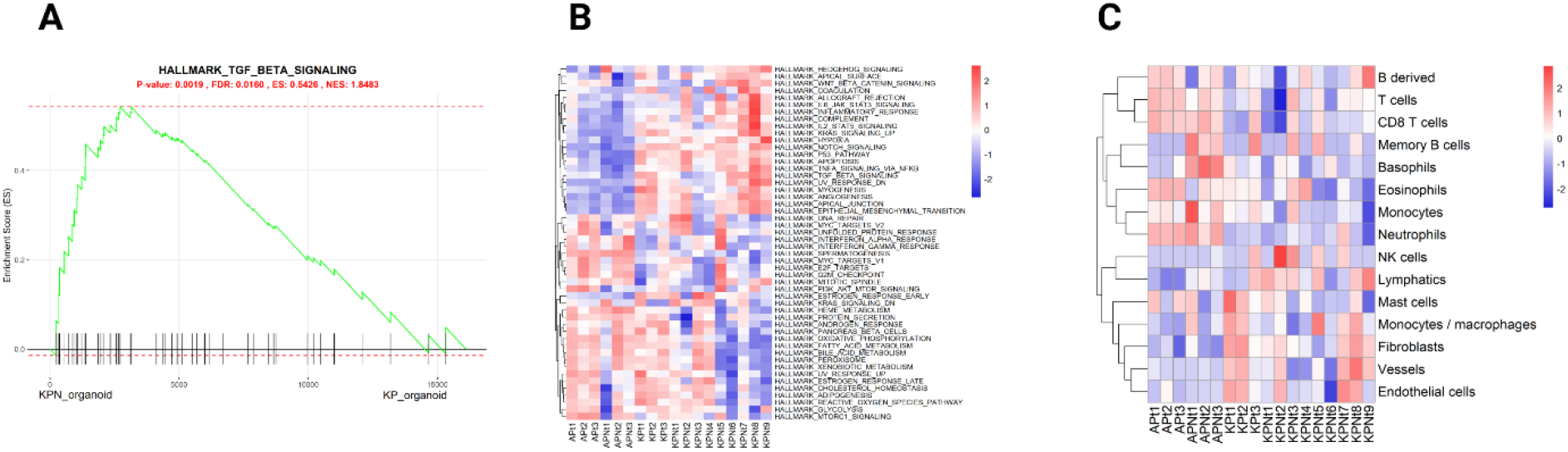
Gene set enrichment analysis and MCP analysis. A) Enrichment plot for TGF-beta signalling Hallmark gene set for KPN vs KP organoid groups with p-value, FDR value, enrichment score (ES) and normalised enrichment scores (NES). The X-axis is all the genes in the data experiment pre-ranked by the metric, where each black bar is the gene in this gene set (pathway) and the Y-axis details the level of enrichment via an Enrichment Score (ES). B) Single sample gene set enrichment analysis for individual samples displayed in a heatmap. C) mMCP analysis with infiltrating cell population estimates visualised in a heatmap.

## Discussion

Every day significant amounts of molecular data are created from biological samples at an ever-reducing cost, shifting the challenge from data acquisition to data analysis and interpretation that can inform deeper understanding of biological processes. Such understanding is essential in order to improve our understanding of disease and identify mechanistic signaling that can aid in diagnosis, prediction of disease outcomes or in the development of new therapeutic strategies (Gambardell, 2020). An example of how important interrogation of molecular data can be is reflected in the worldwide response to the COVID-19 pandemic, where rapid interpretable data underpinned a meaningful mitigation response to the pandemic’s impact on health and society (Lai et al., 2020).

Data analytical pipelines require specific skill sets, such as data informatics and specific programming, which are not currently in the armamentarium of traditional “wet-lab” scientists. An increased focus on biomarker development and target-drug discovery for personalized medicine requires results generated by gene expression profiling to be interrogated using high-performance computing and potentially with advanced Artificial Intelligence (AI) / Machine Learning (ML) algorithms, again requiring the use of complex bioinformatics tools. For the non-computational biologist, MouSR, with its intuitive structure and user-friendly navigation, enables rapid point-and-click publication-ready analysis of highly complex information, where significant volumes of data can be analysed using multiple methodologies on a single app. MouSR provides a unique opportunity for non-specialist users to analyse their data using customised easy-to-use bioinformatic tools, while also having dual functionality embedded within the App to investigate disease-specific models and algorithms that offer deeper insights to facilitate simultaneous classification between human- and mouse-derived data. The user can choose to deploy all the features within our intuitive transcriptional analysis pipeline for comprehensive work-up, or in other instances the user might decide to utilise only a selection of the available options within MouSR for their bespoke analysis requirements.

The application is internet accessible, but by making our source code freely available, MouSR provides an open-source option for individual users or institutes to install their own instance on local computers/servers. Furthermore, given the remarkable growth in the R programming language community, the MouSR tool provides an adaptable template for further development that will is not limited by recurring software fees. As a clear demonstration of the utility of MouSR, utilising our previously published data, we rapidly reproduced a number of the main molecular findings from the original study in a matter of minutes.

During its development, decisions were made to broaden the range of analyses that MouSR could offer, which in turn leads to a number of limitations which we acknowledge. Multiple analysis methods in MouSR have been created using different libraries under different version of R; finding the best R version that can suit them all and at the same time can accommodate the shiny server version and CentOS server can be challenging. For instance, there is a recently published library called “Interactive Complex Heatmap” (Zuguang Gu and Hübschmann, 2021), which provides an easy-to-use tool for constructing highly customizable heatmaps, especially for analysing DESeq2 results. However, based on the version of R and shiny that we are using, we preferred to deploy the “heatmaply” package. It is worth noting that although this single purpose tool can provide highly customizable heatmaps, it does not have the same breath of capabilities or versatility in comparison with MouSR. Additionally, MouSR (as a free, open-source tool), can provide more analysis methods than other existing free analytical apps currently available. For instance, DEApp (Li and rade, 2017) is only focused on differential expression analysis of count-based NGS data.

In addition, TCC-GUI (Wei Su et al., 2019) uses differential expression pipelines with robust normalization and simulation data generation under various conditions, however it does not include gene set enrichment analysis and MCP/mMCP counter analysis. The START tool (Nelson et al., 2017), while having a number of specific functionalities, again does not include gene enrichment analysis and MCP/mMCP counter analysis. Finally, the GENAVi application (Reyes et al., 2019) can provide certain analyses, however does not include MCP/mMCP counter analysis, multi-group comparison or dual functionality for both human and mouse derived data when compared to MouSR. Despite these limitations, the MouSR architecture design provides a structure that offers, in the future, the possibility to implement new types of bespoke analysis pipelines and graphical outputs with precise functionalities within the open-source R programming language, facilitating access to thousands of statistical packages which are continually released and updated globally.

In summary, MouSR is a freely-available tool that provides a user-friendly graphical interface for biological characterisation and interrogation of transcriptional datasets. Approaches such as ours help remove a bottleneck in biological discovery for users with limited programme skills, enabling them to perform statistically-controlled bioinformatics analyses to make valid biologically-informed conclusions more precisely.

## Supporting information

Supplementary File 1 (Input 1)

Supplementary File 2 (Input 2)

Supplementary File 3 (MouSR tutorial))

## Acknowledgments

This project has been developed as a part of the CRUK-funded ACRCelerate colorectal cancer pre-clinical accelerator program. The ACRCelerate project brings together a European-wide consortium of basic and clinical scientists at the forefront of CRC research to interrogate a suite of state-of-the art preclinical models. The overarching aim of ACRCelerate is to generate robust and reproducible preclinical data to de-risk future clinical trials via patient stratification. We acknowledge ‘VolcaNoseR’ for their Volcano plot and ‘DEApp’ for their first page template inspiration.

We thank the collaborative network across Belfast, Oxford and Glasgow for feedback and discussions during development. We also thank the IT department at Queen’s University and Crispin Miller at the Beatson institute for advice and support.

